# Integrating and refining the predictions of ensembled plant effector detection programs using machine learning

**DOI:** 10.64898/2025.12.17.694896

**Authors:** Love Odunlami, Dan MacLean

## Abstract

Effectors are pathogen proteins that facilitate infection by manipulating plant immunity. Computational programs have been developed that identify effectors from sequence data. Many of these programs use internal models that have unavoidable biases due to their training processes and the diverse nature of effector sequence, function and phylogeny. Each programs ability to predict effectors across a broad range of plant pathogens is therefore limited. We hypothesised that a meta-predictor constructed using machine learning (ML) approaches could integrate predictions from multiple programs and improve our ability to predict effectors more accurately from bacteria, fungi, and oomycetes. We trained a range of classifiers using classical ML approaches and deep neural networks (DNNs), then selected eight: Random Forest (RF), Support Vector Machine (SVM), Extreme Gradient Boosting (XGBoost), and five DNNs for evaluation. The training, test and validation data were carefully curated from effector and non-effector annotated sequence derived from the training and sample data of six programs: EffectorP 3.0, deepredeff, WideEffHunter, EffectorO, EffectiveT3, T3SEpp. The models were tested against existing programs on a test dataset, and we observed better performance from our models. The best-performing model was a DNN (Model_2) that balanced improved sensitivity with specificity across the three taxa. We observed using SHAP that all the features contributed to the output of Model_2, which might be the reason for its superior performance. The DNN was developed into a package, *fimep*, to allow easy use of our model.

## Introduction

The projected increase in global human population from the current 8.2 billion to 10.3 billion by 2080 (United Nations 2024) will increase food demand and, to prevent hunger require great improvements in crop production and the overall food system. Losses to plant pathogens are therefore a great threat to food security. Taxonomically diverse pathogens cause huge amounts of crop loss. The fungi *Magnaporthe oryzae* causes an annual yield loss of between 20-30% in rice (Weitao et al. 2019), while *Puccinia striiformis* f. sp. *tritici* (wheat stripe rust) causes loss of up to $5 billion annually (Figueroa, HammondlKosack, and Solomon 2018). Bacterial pathogens cause global losses of over $1 billion annually (Islam et al. 2024), and the oomycete *Phytophthora infestans*, which causes late blight of potato and tomato, resulting in an estimated yield loss of over $10 billion (Dong and Zhou 2022).

Infection is a dynamic process in which pathogens utilise a unique repertoire of effectors that disrupt the complex plant immune system (Dalio et al. 2018; Marín, Uversky, and Ott 2013; Xiang, Stojilkovic, and Gheysen 2025). These effectors are molecules secreted by pathogens to manipulate host physiology by masking damage-or pattern-associated molecular patterns (DAMPs and PAMPs), hijacking immune signalling pathways, and triggering hypersensitive cell death (Dodds and Rathjen 2010; Lo Presti et al. 2015; Toruño, Stergiopoulos, and Coaker 2016). Characterising effectors remains challenging due to their highly diverse sequence and function, rapid evolution, functional redundancy and complexity, and absence of conserved features across different kingdoms (Calia et al. 2024; Jones and Raffaele 2025; Lovelace et al. 2023; Mukhi, Gorenkin, and Banfield 2020; Sperschneider et al. 2015).

With the rise of high-throughput sequencing techniques and artificial intelligence (AI), computational approaches have been developed to predict candidate effector proteins from sequence data. Despite their critical role, effector proteins remain difficult to characterise; early tools relied on predefined features such as protein size, cysteine content, secretion signals, and motif presence (Sperschneider et al. 2016; Stam et al. 2013; Yu et al. 2021; Xue et al. 2019). However, more recently, machine learning (ML) has allowed for the inclusion of multi-dimensional and non-linear features that better capture the diversity of effector sequences (Kristianingsih and MacLean 2021; Nur et al 2023; Sperschneider and Dodds 2022). Effector prediction algorithms using support vector machine (SVM) (Yejun Wang et al. 2014), random forest (RF) (Nur et al 2023; Tang et al 2024), and, more recently, deep learning (DL) models like convolutional neural networks (CNNs) (Kristianingsih and MacLean 2021; Yansu Wang, Luo, and Zou 2022; Xue et al. 2019), recurrent neural networks (RNNs) (Xue et al. 2018), and hybrid CNN-RNN (Jing et al. 2021; Kristianingsih and MacLean 2021) have been employed to learn from raw sequence data, physicochemical profiles, and amino acid k-mer patterns.

The tool EffectorP 3.0 uses an ensemble classifier (Naïve Bayes classifier and Decision tree) to classify fungal or oomycete effectors as either apoplastic or cytoplasmic with high accuracy (Sperschneider and Dodds 2022). WideEffHunter predicts on oomycete and fungal effectors. It does not use ML algorithms instead relying on sequence characteristics including the presence of effector motifs, domains, and homology to validated effectors (Carreón-Anguiano et al. 2022). deepredeff uses an ensemble of deep learning models to predict effector proteins from oomycetes, fungi, and bacteria. For bacteria, deepredeff uses a weighted average of CNN-long short-term memory (LSTM), CNN-gated recurrent unit (GRU), and GRU-Embedding, while the fungal and oomycete effector prediction is based on a single CNN-LSTM model (Kristianingsih and MacLean 2021). EffectorO, an oomycete effector classification tool, uses a random forest classifier and has shown improved performance in predicting effectors even in the absence of conserved motifs (Nur et al 2023). EffectiveT3 is a Type III secreted effector prediction tool that uses a Naïve Bayesian classifier to recognise N-terminal signal peptides (Arnold et al. 2009). T3Sepp is another Type III secreted effector prediction tool that combines multiple homology-based methods with ML models. It uses a linear model that weights the prediction scores from homology-based screening with a Markov model, SVM, decision tree, and CNN-LSTM to improve the overall accuracy and reduce the high false-positive rate (Hui et al. 2020). While these approaches have improved effector prediction within specific kingdoms, their performance often varies across taxa, and many fail to generalise beyond their training sets and close taxa despite claims of accuracy over wider domains like kingdoms. Moreover, the biological and algorithmic diversity of different effector prediction programs frequently results in varied predictions with limited overlap; Nur et al (2023) highlighted a 42% overlap between EffectorP 3.0 and EffectorO.

We hypothesised that integrating predictions from multiple kingdom-specific effector prediction programs and using these collectively as features to train ML or DL algorithms would yield accurate and balanced effector classification across pathogen kingdoms. We trained RF, SVM, XGBoost and DNN algorithms on prediction output from six effector prediction programs on experimentally verified effectors and non-effectors and showed that the models had better performance with better balanced sensitivity and specificity than any individual program.

## Methodology

### Data curation and encoding

We curated a unique set of 1458 effectors and 2214 non-effectors from the training datasets provided by: EffectorP 3.0 (Sperschneider and Dodds 2022), deepredeff-0.1.1 (Kristianingsih and MacLean 2021), WideEffHunter-1.0 (Carreón-Anguiano et al. 2022), EffectorO (Nur et al 2023), EffectiveT3-1.0.1 (Arnold et al. 2009), and T3SEpp (Hui et al. 2020). We created a balanced negative-to-positive training (999), test (n=1293), and validation (n=1290) dataset using the scikit-learn-1.5.1 library (Pedregosa et al. 2011), then one-hot encoded the prediction output from each program as 1 for effectors and 0 for non-effectors using the pandas-2.23 library (McKinney et al. 2011) in Python-3.11.8. The balanced dataset ensured that for each kingdom (Bacteria, Fungi, and Oomycete), the number of effector and non-effector were equal across all three datasets (additional Table S1).

### Training, hyperparameter optimisation, and model selection

We trained one each of RF, SVM, and XGBoost based models and five DNNs. For the classical ML models, we evaluated and optimised them using a GridSearchCV and cross-validation (CV), while the DNNs were optimised using RandomizedSearchCV with CV. To reduce overfitting and compare the models, we monitored the accuracy, sensitivity, specificity, and F1 score. RF and SVM were used from the scikit-learn-1.5.1 library, while XGBoost and the DNNs were from the XGBoost (Chen and Guestrin 2016) and Keras (Chollet et al. 2018) libraries, respectively.

We evaluated the models on the test and validation data, compared their performance using Hamming distance and selected the best-performing models based on their accuracy, sensitivity, specificity, and F1 score. The Hamming distance and confusion matrix were computed using pairwise distance function and confusion matrix function from scikit-learn-1.5.1 library respectively. Plots were created using ggplot2 within the tidyverse-2.0.0 package (Wickham et al. 2019). To understand the contribution of each feature to observed prediction, we generated a beeswarm summary plot using the SHAP-0.47.2 package (Lundberg, Erion, and Lee 2018).

### Ensemble models

To create an ensemble of the models, we used two techniques: weighted average and maximum voting.

The weighted average was calculated using:

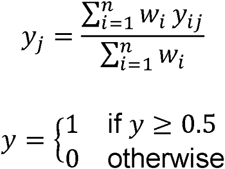

The majority voting was estimated using:

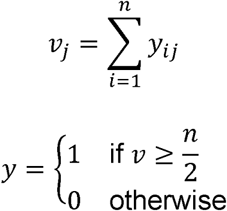

Where y_j_ is the weighted average prediction for sample j, w_i_ is the weight, y_ij_ is the prediction value for sample j of the i^th^ model, and n is the total number of models, ν_j_ is the number of models that voted 1 for the sample j.

The workflow from data acquisition to effector prediction using trained models is shown in Figure 1.

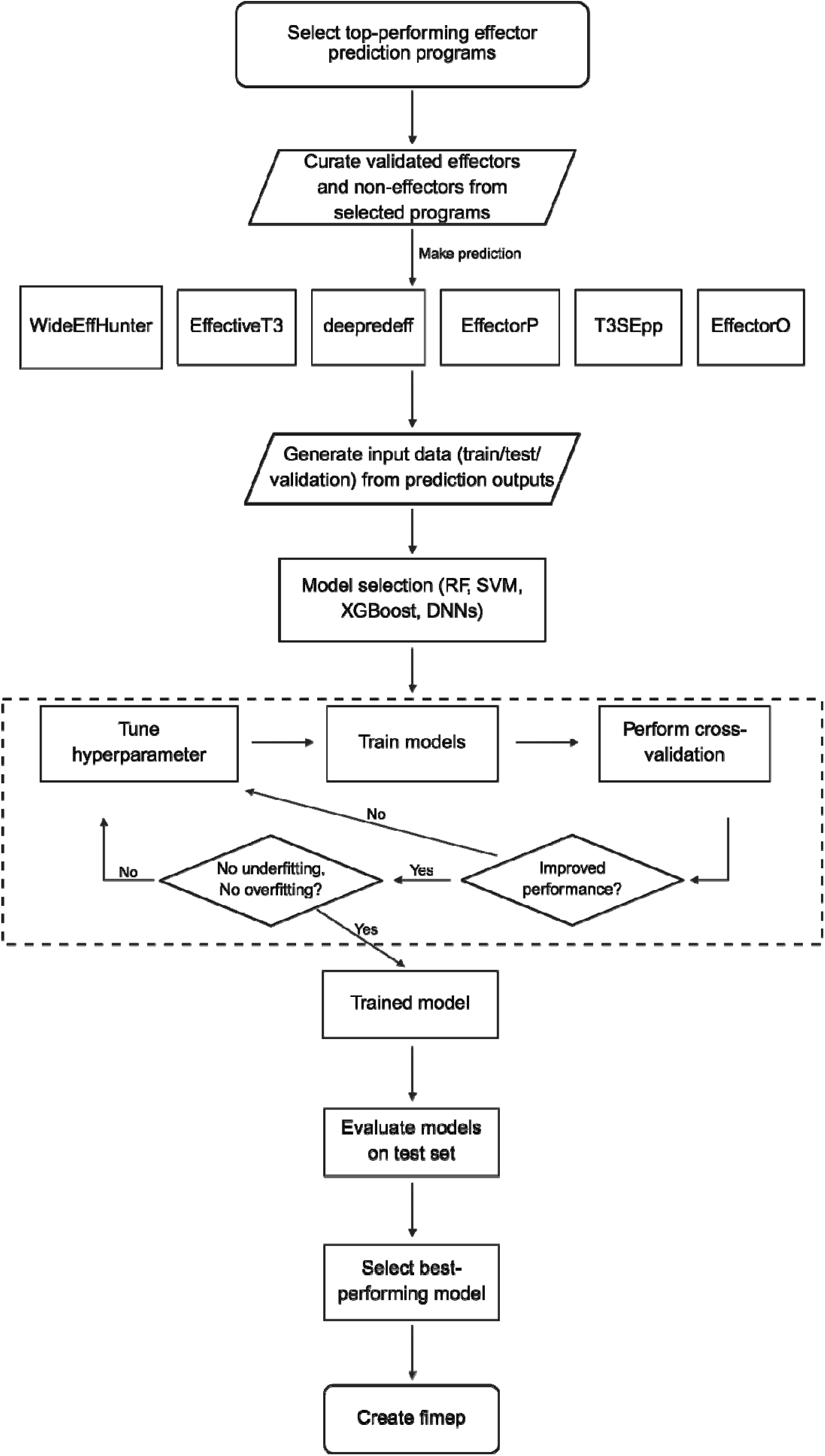

### Evaluation metrics

We evaluated the performance of the models using the metrics explained below. Accuracy: the proportion of predicted effectors and non-effectors to the total prediction.

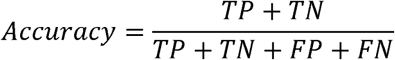

Sensitivity: the ratio of the correctly predicted effectors to all the effectors.

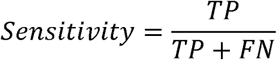

Specificity: the proportion of the correctly predicted non-effectors to all the non-effectors.

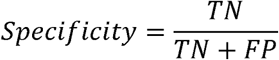

F1 score: the harmonic mean of the precision and recall (sensitivity).

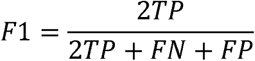

## Data availability

The data for this study, compiled from predictions across three taxa (fungi, bacteria, and oomycetes), are presented in Table 1. The programs make predictions based on sequences from the taxonomic groups indicated. The data used in this study, the models developed, and all the scripts are available on GitHub at https://github.com/LoveBio/Effector_prediction.

**Table 1:**
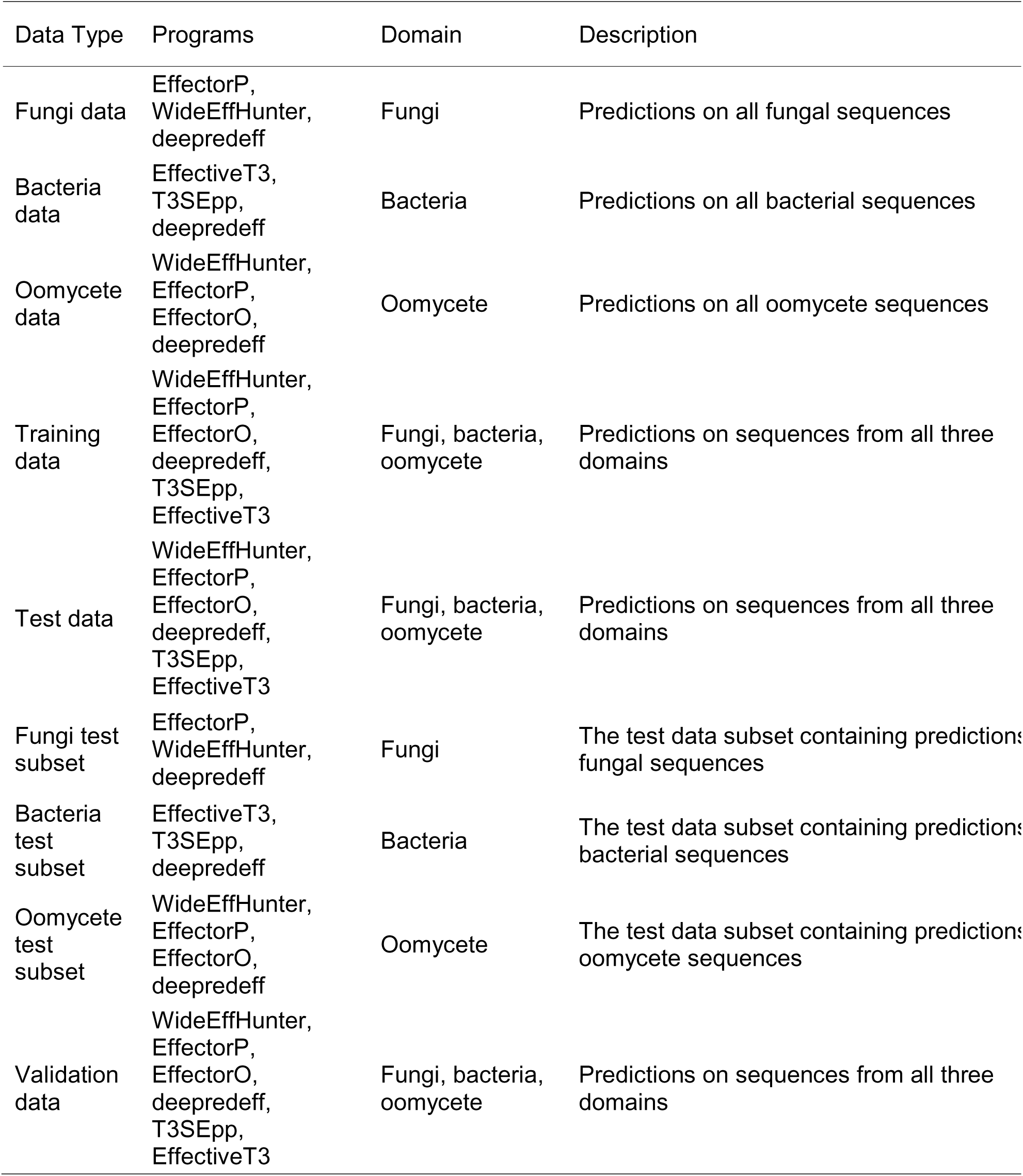
Overview of the datasets and prediction programs used in this study.

### Software implementation

We developed a command-line package called *fimep* (for integration of multiple effector predictions) that packages the top-performing DNN model, enabling easy integration into any effector prediction pipeline. The package, along with installation instructions, is available on GitHub (https://github.com/LoveBio/fimep) and the Python Package Index (PyPI) at https://pypi.org/project/fimep/. *fimep* includes modules for preprocessing outputs from all programs used in this study, encoding and formatting input data, and generating more robust and accurate overall predictions.

## Results

### Program identification

We conducted a literature review to identify standalone effector prediction programs that are accessible, functional, supported, trained on experimentally validated effectors, and with publicly available training and validation datasets. We selected EffectorP and WideEffHunter that predict on fungi and oomycetes pathogens (Sperschneider and Dodds, 2022, Carreón-Anguiano et al., 2022), deepredeff predicting on fungi, oomycetes, and bacteria pathogens using separate models (Kristianingsih and MacLean, 2021), T3SEpp and EffectiveT3 predict on bacteria pathogens (Arnold et al., 2009, Hui et al., 2020), and EffectorO (Nur et al., 2023), which predicts on oomycete pathogens. Since the start of this project, EffectiveT3 has been moved to the Life Science computer Cluster (LiSC) Galaxy (version 2.0.2) and the original version, wrapped as a web-based tool (Cock et al. 2013).

### Effector prediction programs give overlapping but distinct results on the same input data

To test the hypothesis that programs generate non-overlapping predictions, we generated predictions across three distinct datasets: bacteria, fungi and oomycete using the process described in the methods. The combined outputs of all programs are shown as a heatmap in Figure 2.

**Figure 2:**
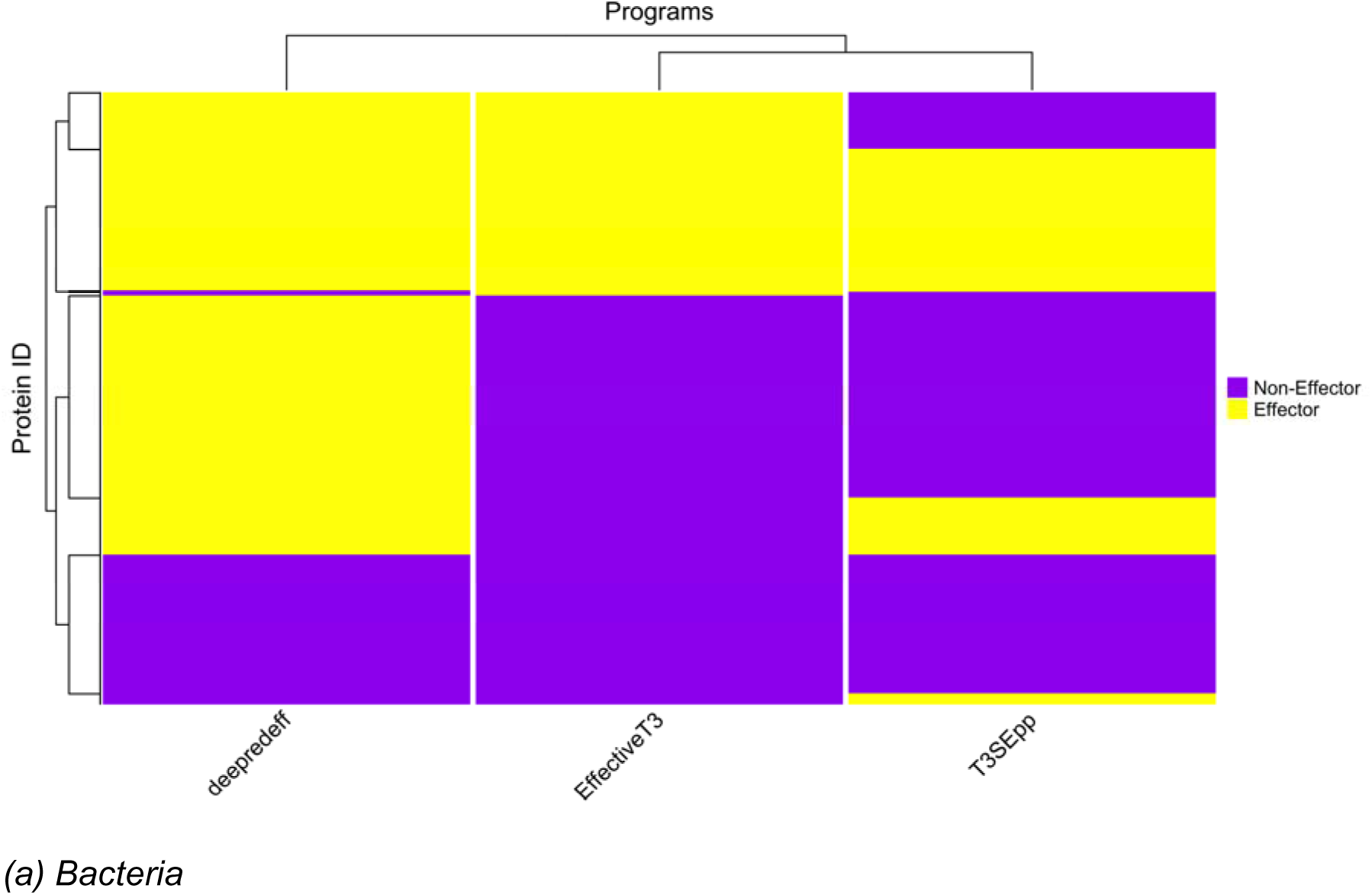
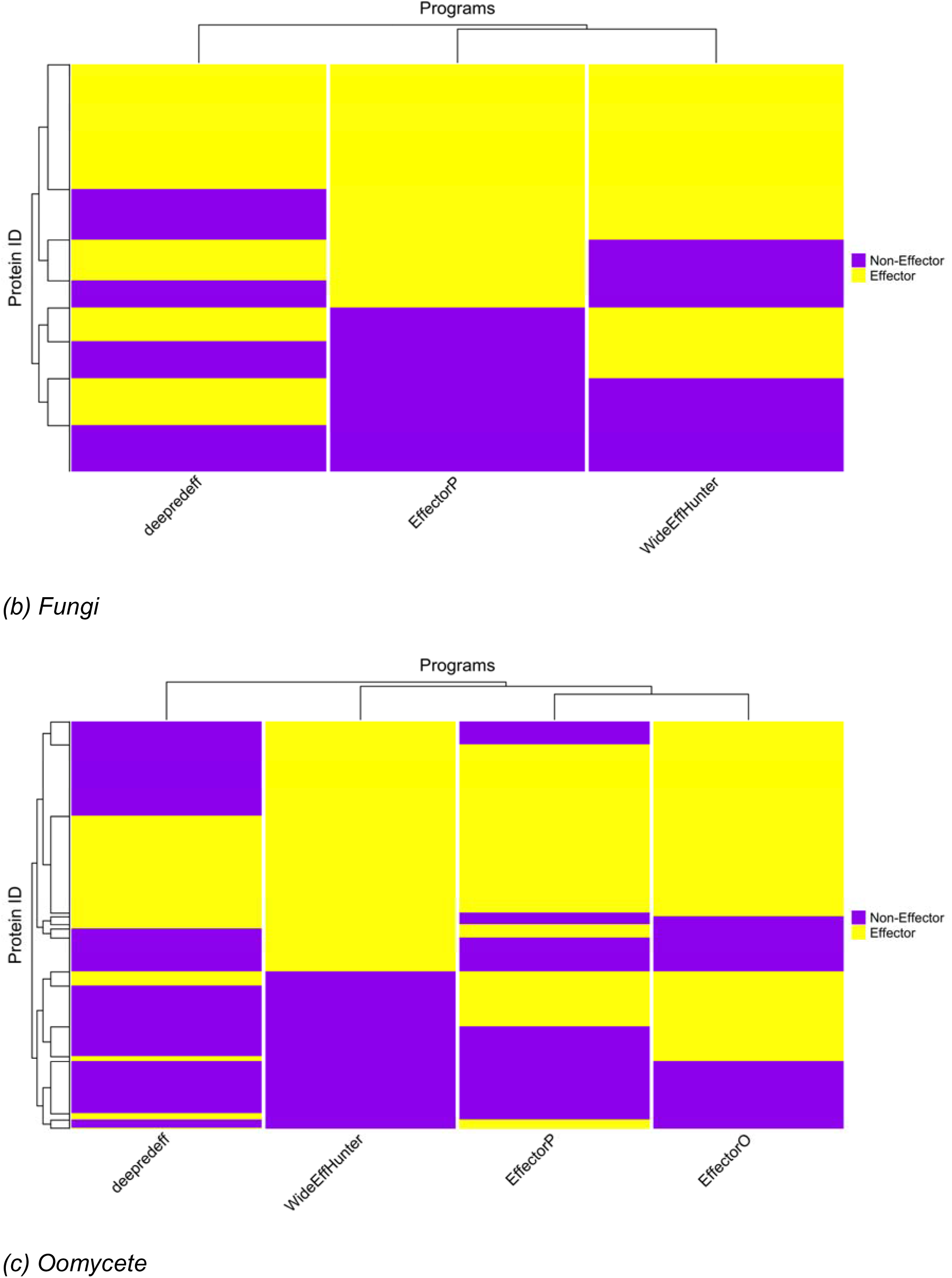
Clustered heatmap of effector protein predictions from kingdom-specific programs. **(A)** Prediction patterns from deepredeff, EffectiveT3, and T3SEpp on bacterial data **(B)** Prediction on Fungal data using deepredeff, EffectorP, and WideEffHunter **(C)** Prediction on oomycetes data using deepredeff, EffectorO, EffectorP, and WideEffHunter

For bacterial proteins (Figure 2 (a)), we observed that 45.84% of the proteins were predicted by all the programs. Each program demonstrated a significant number of unique predictions, indicating specialised capabilities. deepredeff had the highest unique prediction rate at 33.25%, followed by T3SEpp at 11.02%, and EffectiveT3 at 9.89%. Additionally, about 18.73% were predicted by a combination of two, but not all, programs.

The prediction on the fungal data (Figure 2 (b)), showed a similar pattern of varied predictions. While 36.59% of the proteins were predicted by all the programs, a large percentage were unique to individual programs. deepredeff uniquely predicted 24.03% of the proteins, WideEffHunter uniquely predicted 19.04%, and EffectorP uniquely predicted 14.97%. We also found that 30.78% were predicted by a subset of any two of the programs.

The prediction on the oomycete (Figure 2 (c)) further depicts the predictive diversity among the programs. As with the other dataset, there was an intersection where all four programs predicted 36.59% of the proteins similarly. We noted a greater distribution of predictions across subsets of programs, with 22.97% of proteins being predicted by three programs and 21.24% by two programs. The unique predictions varied significantly: deepredeff had the highest unique rate at 19.11%, followed by WideEffHunter (11.79%), EffectorO (8.33%), and EffectorP (2.95%).

The heterogeneity observed across all three datasets supports the idea that there is a potential approach to improve effector prediction involving training a meta-model that uses these individual strengths, potentially combining the unique predictions of each program and possibly creating a more robust set of predictions.

### Model training and selection

We aimed to create a model that would integrate outputs from all these programs and make a meta-prediction from their initial predictions. We reasoned that this would be a low volume dataset, with number of columns as per the number of programs used and the number of rows as per the number of effectors and non-effectors in the shared training sets. Hence it seemed that classical ML algorithms would be suited as well as currently more popular DNN based methods, in small sizes. We selected a range of likely algorithms and began training. We performed a grid hyperparameter search with fourfold cross-validation for the classical ML algorithm (RF, SVM, and XGBoost) and a randomised hyperparameter search with fivefold cross-validation for the DNN (Model_1 to Model_5) to identify classifiers with high accuracy. The values of the hyperparameters are listed in additional Table S2. We trained a total of 857,000 models and selected the eight best-performing models based on their ranking, as determined by accuracy, precision, recall, and F1 score (Table 2). An evaluation of the validation data (Figure 3) showed that all the models maintained a high evaluation metric. Some DNN models, such as Model_1 and Model_4, had a slightly different pattern, with high sensitivity (90.17% and 90.38%, respectively) and relatively low specificity (74.48% for both).

**Figure 3:**
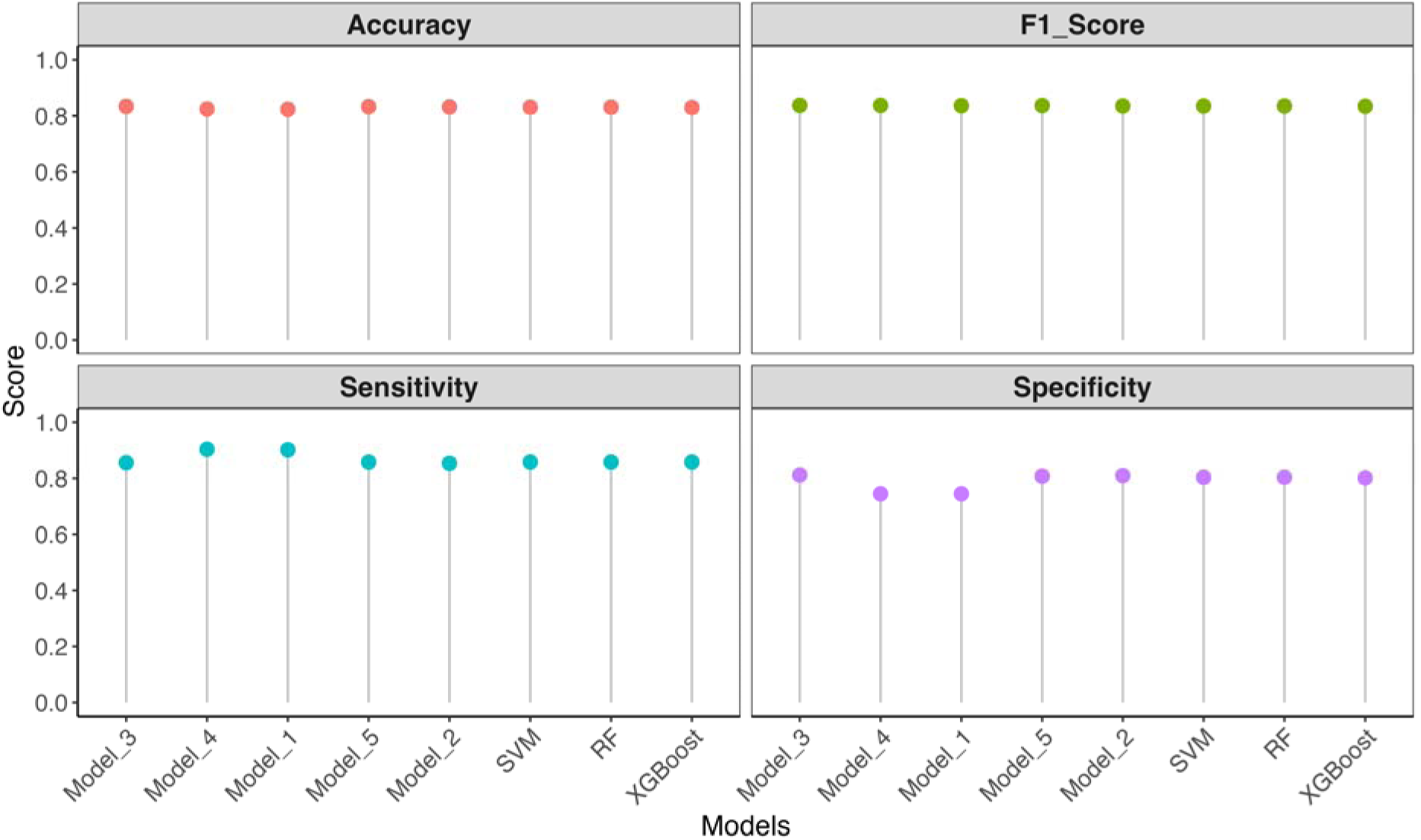
Comparison of selected models on the validation data. This plot shows a direct comparison of the models’ ability to balance sensitivity and specificity while maintaining high overall accuracy. Models were ranked by F1 score. XGBoost, extreme gradient boosting; RF, Random Forest; SVM, Support Vector Machine; deep neural network models (Model_1 to Model_5)

**Table 2:** Cross-validated performance of selected models across accuracy, precision, recall, and F1 score.

| Model<br>(Cross-Validation) | Mean<br>Accuracy (%) | Mean<br>Precision (%) | Mean<br>Recall (%) | Mean F1<br>Score (%) |
| --- | --- | --- | --- | --- |
| RF (4) | 85.98 | 84.48 | 88.37 | 86.28 |
| SVM (4) | 85.78 | 84.08 | 88.57 | 86.14 |
| XGBoost (4) | 85.38 | 84.11 | 87.56 | 85.65 |
| Model_1 (5) | 86.28 | 84.49 | 88.97 | 86.60 |
| Model_2 (5) | 86.08 | 84.56 | 88.37 | 86.38 |
| Model_3 (5) | 86.08 | 84.53 | 88.36 | 86.35 |
| Model_4 (5) | 85.88 | 84.36 | 88.17 | 86.18 |
| Model_5 (5) | 85.78 | 84.27 | 87.97 | 86.06 |

### Model performance and comparison

We compared the predictive ability of the selected models on the test data with the goal of finding the top performer and understanding the similarity and differences in how each model makes its prediction.

Among the classical ML models (Figure 4 (a)), XGBoost showed a slight performance edge, correctly identifying 385 (80.00%) proteins as non-effectors and 441 (91.70%) as effectors. Its performance was marginally better than RF and SVM, which both correctly classified 440 (91.50%) effectors and 384 (79.80%) non-effectors. For the DNN (Figure 4 (b)), Model_2 and Model_4 were the best. Model_4 excelled at correctly classifying 449 (93.3% vs 91.50%) effectors, while Model_2 correctly identified higher 388 (80.70% vs 74.80%) non-effectors. Model_1 and Model_4 showed similar classification for effectors (449 – 93.30% each), though Model_4 correctly identified an additional five non-effectors (a 1.1% increase). Model_3 and Model_5 mirrored each other, correctly predicting 386 (80.20%) non-effectors, but Model_5 predicted more effectors than Model_3 (440 vs 438). Similarly, Model_2 and Model_5 correctly predicted 440 (91.50%) effectors and a close prediction number for non-effectors (388 vs 386).

**Figure 4:**
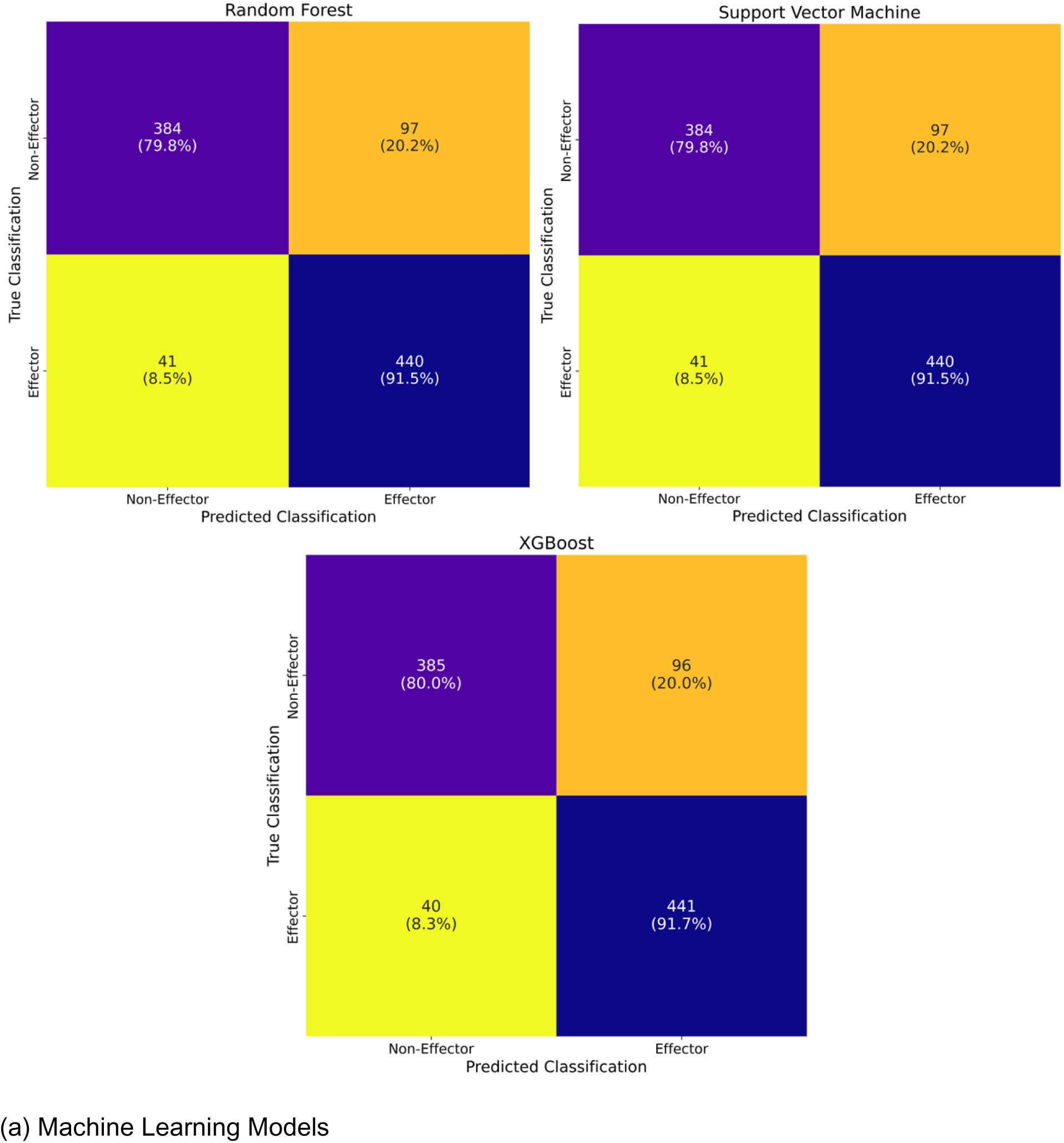
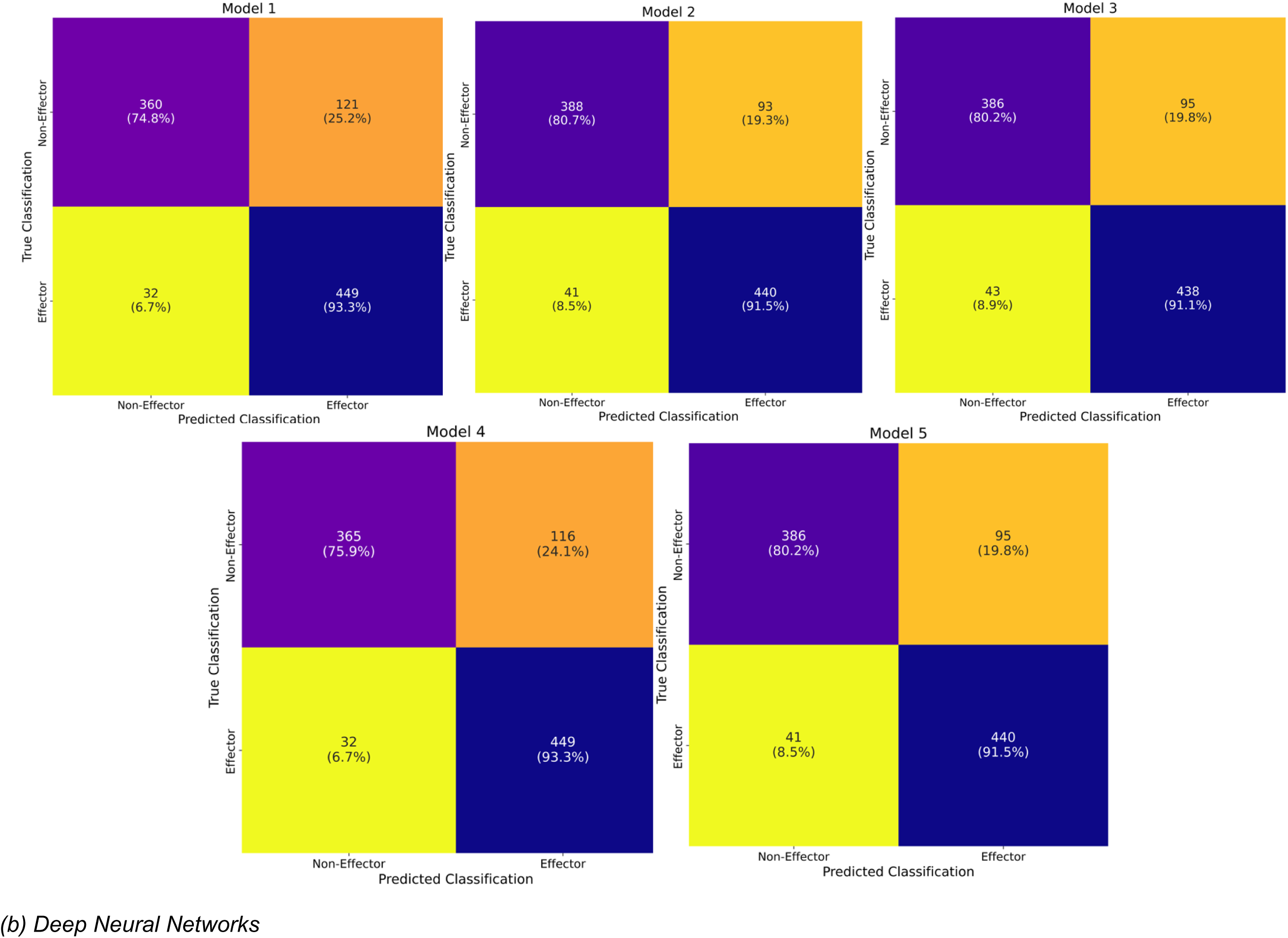
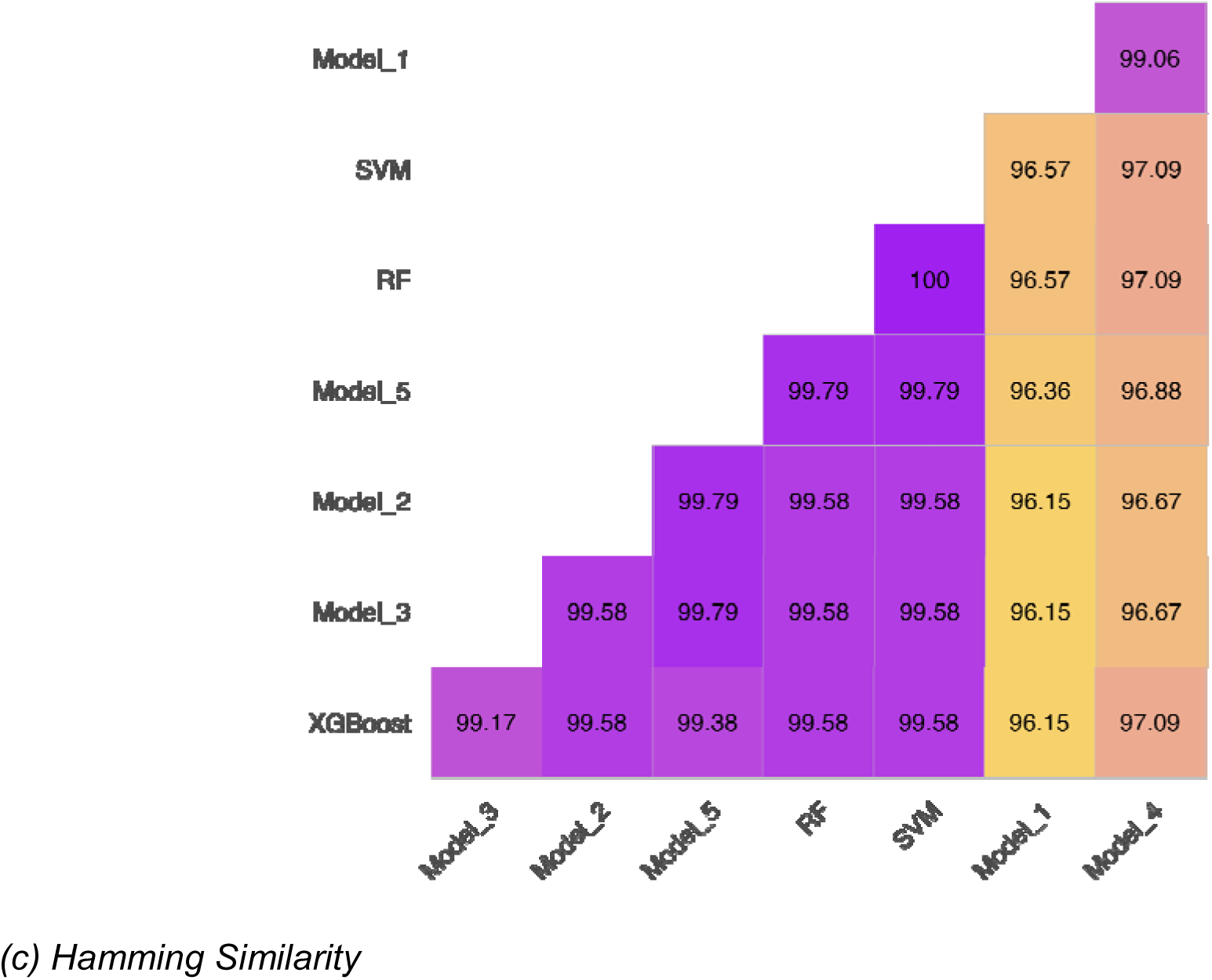
Classification performance of all models on test data, with confusion matrices and Hamming similarity analysis. **(A)** Confusion matrices comparing the predicted labels to the actual labels for the machine learning models (XGBoost, RF, and SVM). **(B)** Confusion matrices comparing the predicted labels to the actual labels for deep neural network models (Model_1 to Model_5). **(C)** Pairwise Hamming similarity comparing exact prediction matches across all models. XGBoost, extreme gradient boosting; RF, Random Forest; SVM, Support Vector Machine; deep neural network models (Model_1 to Model_5)

To understand the consistency of these predictions, we calculated the Hamming distance, which measures agreement in exact predictions for any pair of models (Figure 4 (c)). The models showed a high degree of agreement, with prediction similarity ranging from 96.15% to 100%. The classical ML models were more similar to each other than they were to the DNNs, ranging between 99.58 and 100%. RF and SVM produced identical outputs, suggesting they could be used interchangeably. Among the DNNs, Model_5 and Model_3 were nearly identical (99.79%), while Model_2, Model_3, and Model_5 showed similarity levels ranging from 99.58% to 99.79%. Model_1 and Model_4 were most similar (99.06%).

All models achieved high performance metrics (Figure 5), confirming their ability to generalise to new, unseen data (Figure S1). To assess overfitting, we inspected learning curves (Figure S1): for the DNN, training and validation loss decreases and then stabilised over epochs, and for the machine learning models, validation and training accuracy increased as stabilise as the size of the training set increased. Sensitivity ranged from 91.06 to 93.34%, specificity from 74.84 to 80.67%, accuracy from 84.10 to 86.07%, and F1 score from 85.44 to 86.79%. Model_2 emerged as the best-performing model, correctly classifying effector and non-effector proteins.

**Figure 5:**
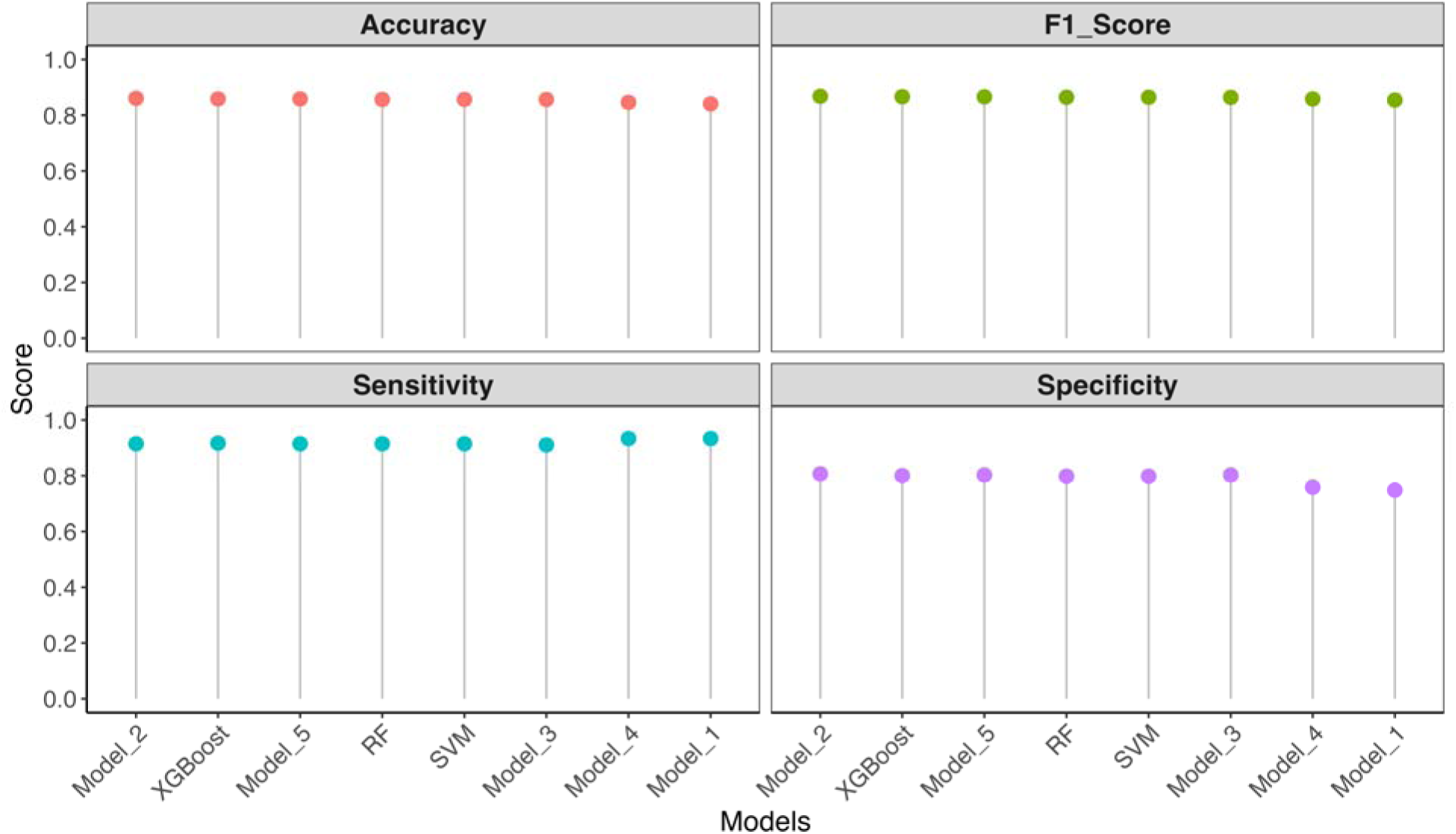
Comparison of model performance based on accuracy, F1 score, sensitivity, and specificity. This plot shows a direct comparison of each model’s ability to balance sensitivity and specificity while maintaining high overall accuracy. Models are ranked by their F1 score, with Model_2 demonstrating the most robust and consistent predictive performance among all models. XGBoost, extreme gradient boosting; RF, Random Forest; SVM, Support Vector Machine; deep neural network models (Model_1 to Model_5)

Its improved specificity and sensitivity, combined with the highest F1 score, established it as the most effective model, followed closely by XGBoost.

### Model Comparisons – Integrated models are more accurate than the effector programs alone

We hypothesised that integrating the predictions from programs as features for the trained ML and DNN models would improve effector prediction accuracy. We aimed to maximise each program’s strengths while minimising its weaknesses, through the models ability to learn patterns, thereby improving effector prediction beyond any single program across kingdoms.

On the oomycete test subset data in Figure 6 (a), the trained models outperformed existing programs. The programs showed an increase in sensitivity and a simultaneous decrease in specificity, resulting in very low F1 scores (35.71 – 71.43%). For instance, deepredeff had the lowest sensitivity (67.14%) but had a high sensitivity (71.43%), while EffectorP, EffectorO, and WideEffHunter had a reversed trend, reaching high sensitivity (92.86%, 94.29%, and 97.14%, respectively) with very low specificity (45.71%, 35.71%, and 50.00%). In comparison, our models achieved high sensitivity, ranging from 90% to 97.14%, while also maintaining specificity (between 51.42% and 71.43%). The models had a higher F1 score, with the top performer achieving 83.87%, significantly exceeding the best program’s F1 score of 78.61%. XGBoost performed exceptionally among the classical ML models (sensitivity: 92.86%, specificity: 68.57%, F1 score: 82.80%). Among the DNN models, Model_2 demonstrated superior performance across all metrics with sensitivity of 92.86%, specificity of 71.43%, and achieving the highest F1 score and accuracy (83.87% and 82.14%, respectively).

**Figure 6:**
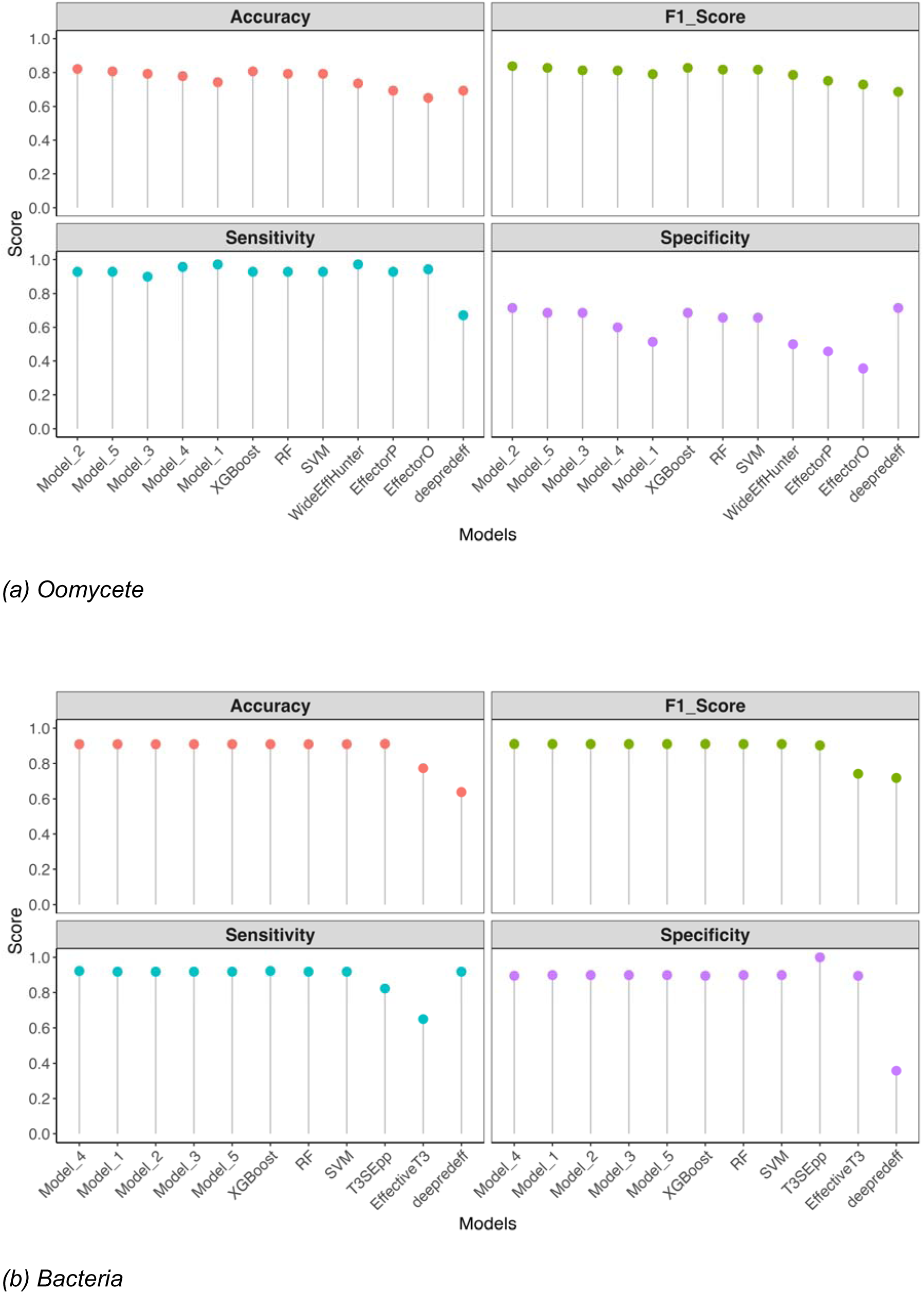
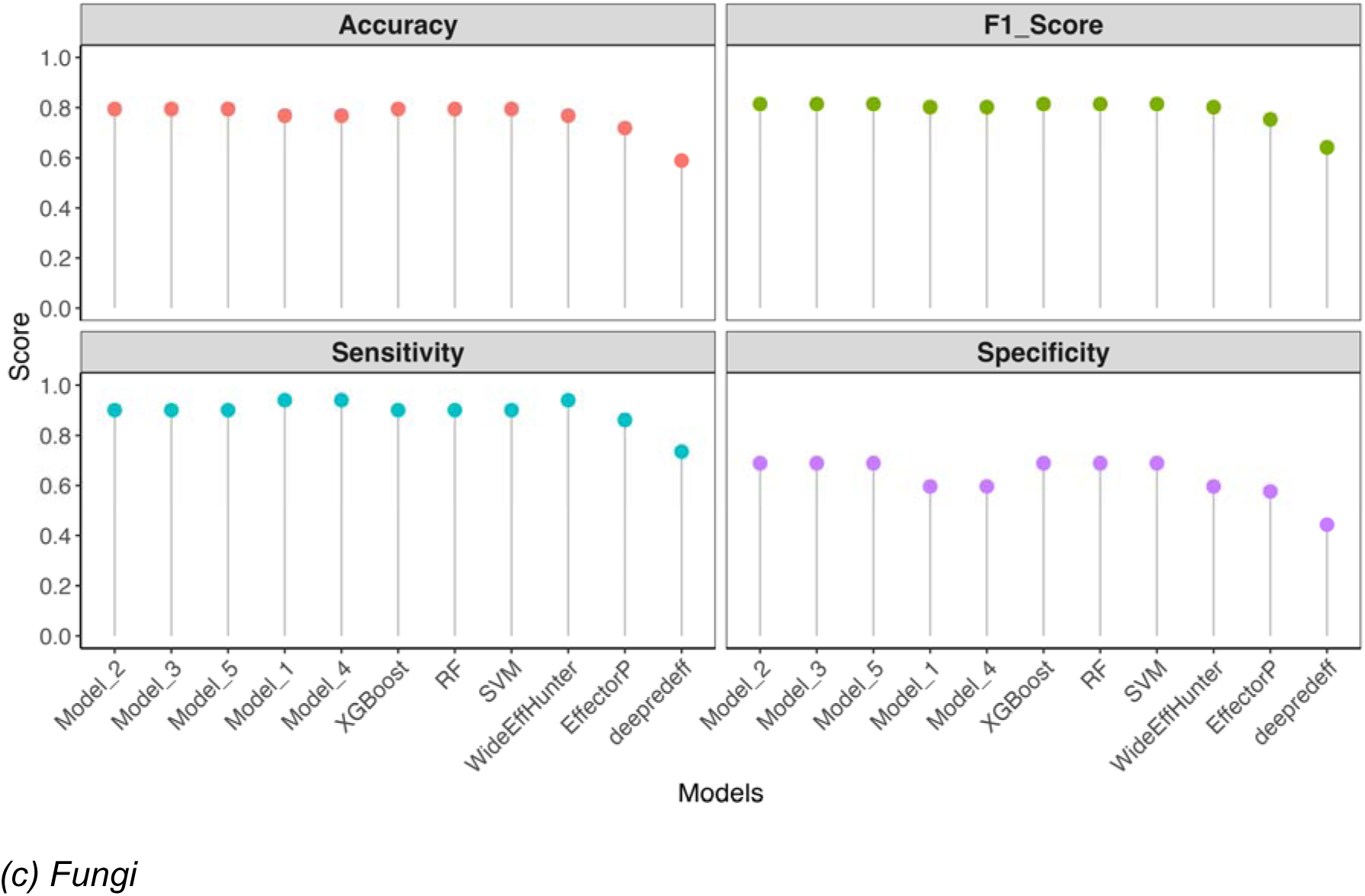
Comparisons of the performance of trained models to the existing programs based on accuracy, F1 score, sensitivity, and specificity. **(A)** Performance comparison using oomycete test subset data, with Model_2 ranked highest among the trained models. **(B)** Comparison using bacterial test subset data, where Model_4 achieved the best performance, though the other models showed closely comparable results. **(C)** The performance of the models and programs on the fungal test subset data showed that Model_2 outperformed other models, although most models achieved comparable performance. Models and programs are first grouped by model type (first the deep neural networks, then the classical machine learning models, and finally the existing programs) before ranking by the F1 score. Overall, Model_2 demonstrates a robust and consistent predictive performance across all taxa. XGBoost, extreme gradient boosting; RF, Random Forest; SVM, Support Vector Machine; the deep neural network models (Model_1 to Model_5)

Evaluating the models on the bacteria test subset data (Figure 6 (b)) showed a superior performance in sensitivity and specificity compared to the programs. deepredeff had high sensitivity (91.93%) and very low specificity (35.77%), while the opposite was observed for EffectiveT3 (sensitivity: 65.00%, specificity: 89.62%). While T3SEpp stood out with high sensitivity (82.31%) and perfect specificity (100%), all our models had better sensitivity and comparable specificity than T3SEpp. XGBoost and Model_4 had the best performance, achieving an F1 score of 91.08%, with all other models performing nearly identically (91.05%). These demonstrate the model’s effectiveness in predicting bacterial proteins.

In predicting the fungi test subset data (Figure 6 (c)), the programs had low specificity (between 44.37% and 59.60%) and high sensitivity between 73.51% and 94.04%. Our models followed a similar trend but consistently achieved a higher sensitivity and a better specificity than the programs. Model_4 and Model_1 had the same performance as the best existing program on our data, WideEffHunter. The classical ML models RF, SVM, and XGBoost, along with the DNN models Model_2, Model_3, and Model_5, were the best-performing models with identical metrics (sensitivity: 90.07%, specificity: 68.87%, accuracy:79.47%, and F1 score: 81.44%).

Our results strongly supported the hypothesis that the trained models integrated individual programs’ predictions (Figure S2), maximising their strength with improved accuracy across all tested kingdoms. The information from the similarity plot and confusion matrix confirmed that the models make predictions with slight differences, suggesting a possibility to ensemble models for a more accurate prediction.

### Comparison of the trained models to an ensemble of the trained models

To test the hypothesis that an ensemble of the models would give a better result than already seen, we developed ensemble models using maximum voting (MVote) and weighted average (WAve). we selected XGBoost, RF over SVM (because they had 100% identical predictions), as well as all the DNNs. The models developed using the maximum voting technique had the lowest accuracy and F1 score (Figure 7) compared to those developed using the weighted average. The MVote models with only the DNN (E_DNN_MVote) had higher accuracy (85.86%) and F1 score (86.61%) compared to models with a combination of the ML and DNN models (E_DNN_ML_MVote), with accuracy and F1 score of 85.65% and 86.44%, respectively. The two ensemble models developed using WAve (E_DNN_ML_WAve; deep neural network and machine learning models, and E_DNN_WAve; only the deep neural network) had identical predictions as Model_2 (accuracy: 86.07%, F1 score: 86.79%). We found that Model_2 contributed the highest weight to the WAve models after a grid search. Therefore, the results from these techniques imply that the ensemble method does not offer any added advantage.

**Figure 7:**
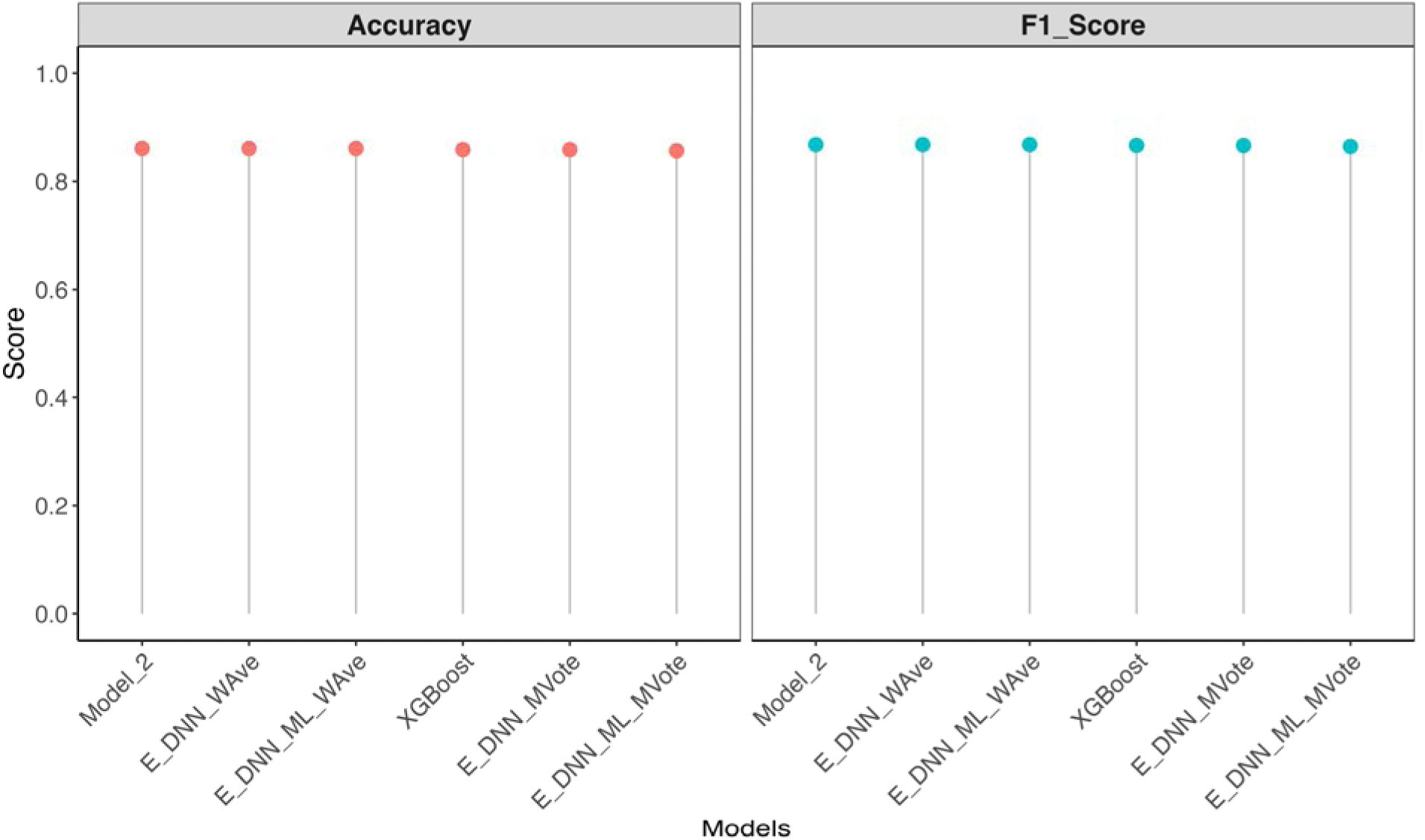
Comparison of the performance of ensemble models developed using maximum voting and weighted average techniques, evaluated by accuracy, F1 score, sensitivity, and specificity. Models were ranked by F1 score. Ensemble models built using maximum voting generally showed lower performance compared to those developed with weighted averaging. However, neither ensemble approach outperformed Model_2, which remained the top-performing model. XGBoost, Extreme Gradient Boosting; the deep neural network models Model_2; E_DNN_WAve, ensemble deep neural network models using weighted average; E_DNN_ML_WAve, ensemble deep neural network and machine learning models using weighted average; E_DNN_MVote, ensemble deep neural network models using maximum voting; E_DNN_ML_MVote, ensemble deep neural network and machine learning models using maximum voting

### Feature contribution pattern differs across machine learning and deep neural network models

We used SHAP (SHapley Additive exPlanations) to understand how features contributed to the prediction outputs of our best models, XGBoost and Model_2. Our analysis showed that each model considered either the WideEffHunter_Non-Effector or T3Sepp_Effector feature most important, but their remaining feature importance and variability differed significantly (Figure 8).

**Figure 8:**
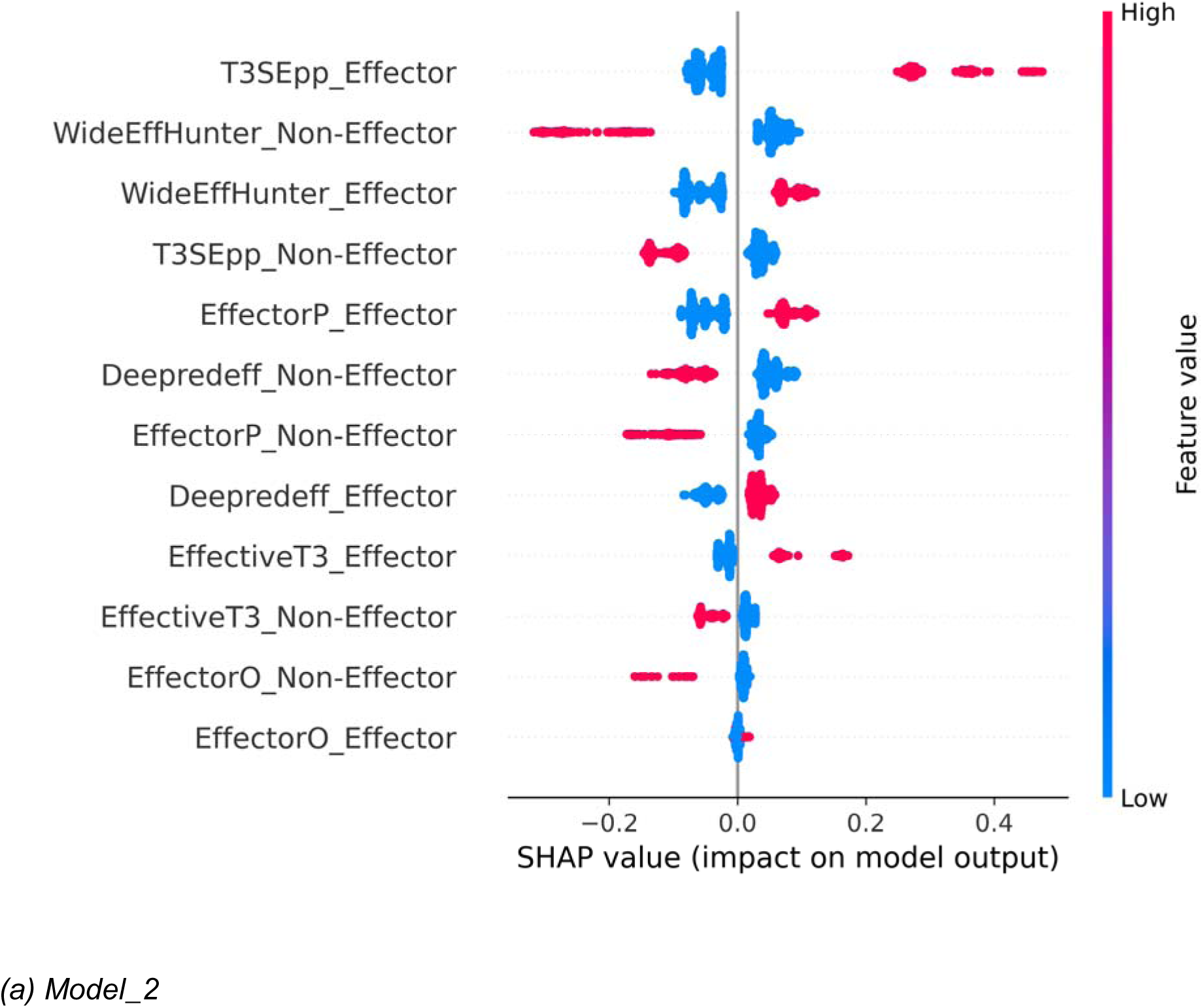
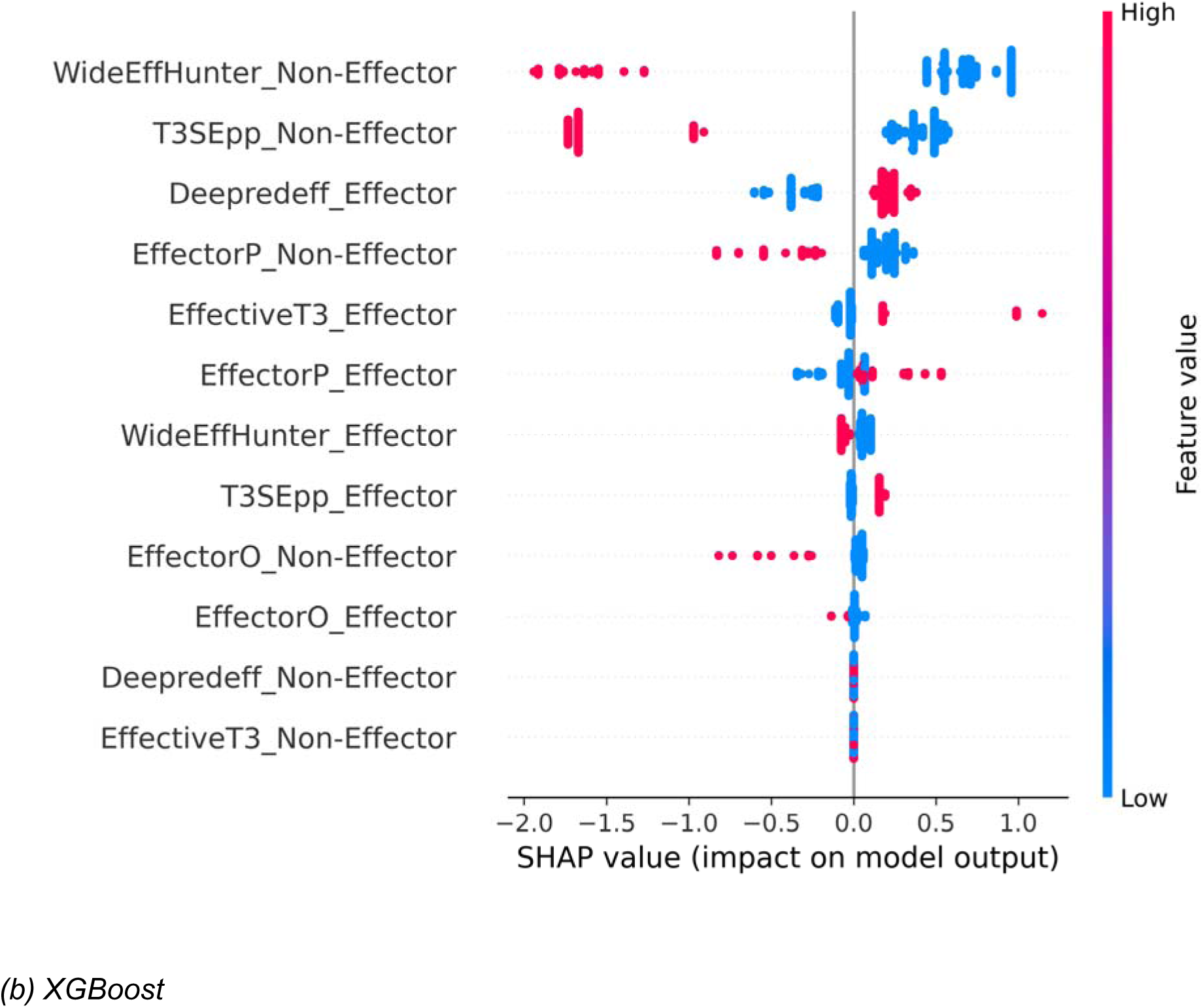
Feature impact on effector prediction models, Model_2 and XGBoost. The y-axis represents the ranked performance contribution of the features to the output, with the most variable features at the top and the least variable features below. The x-axis shows the SHAP values, which quantify each feature’s contribution to the model’s prediction for individual samples. Positive SHAP values indicate that features push the prediction towards classifying as an Effector, while negative values push towards Non-Effector predictions

In Model_2 (Figure 8 (a)), all features contributed to the prediction, with only the EffectorO_Effector feature making a marginal contribution. In contrast, the XGBoost model (Figure 8 (b)) received no contribution from two features: Deepredeff_Non-Effector and EffectiveT3_Non-Effector. A key difference between the two models was the relationship between feature values and their impact on the WideEffHunter_Effector and EffectorO_Effector features. For XGBoost, higher values for the WideEffHunter_Effector feature and EffectorO_Effector consistently had a negative impact on the outcome, which was the opposite of what we observed in Model_2. This lack of contribution from two features and the difference in how the WideEffHunter_Effector and EffectorO_Effector feature influenced the outcome likely explain why XGBoost performed slightly worse than Model_2.

## Discussion

Our study demonstrates the effectiveness of using diverse machine learning and deep learning approaches for predicting effector proteins across multiple kingdoms, demonstrating that integration of prediction from multiple programs significantly improves performance over an individual program. By combining the predictions from six state-of-the-art effector prediction programs including kingdom-specific programs like EffectiveT3 and T3Sepp for bacteria, EffectorO for oomycetes, and multi-kingdom predictors such as EffectorP and WideEffHunter for fungi and oomycetes, and deepredeff for all three kingdoms (Arnold et al. 2009; Carreón-Anguiano et al. 2022; Hui et al. 2020; Kristianingsih and MacLean 2021; Nur et al 2023; Sperschneider and Dodds 2022), we addressed the limitation associated with kingdom specificity, model bias, and accuracy of prediction. While the standalone version of EffectiveT3 became inaccessible during the study, we confirmed that a functional web-based version remains available on the LiSC Galaxy platform. All programs used in this study remain accessible to researchers.

Our result reveal that prediction heterogeneity among programs (Figure 2) can be exploited, and this diversity enabled our models to capture complementary strengths when trained on ensemble outputs, which Tong and Li (2025) showed in their work that an ensemble of base learners outperformed any single model in predicting learning achievements. Notably, Model_2, a DNN, had the best performance, maintaining high specificity and sensitivity while achieving an optimal F1 score on the test and independent validation data.

Comparing our trained models to existing programs revealed the consistent improvement in performance. For oomycete, bacteria, and fungi predictions, existing programs showed striking variation in performance across sensitivity, specificity, and F1 scores. Although our models also showed similar variation in performance for fungi and oomycete predictions, their overall performance was still better than the programs. Our models prediction on bacteria dataset gave more accurate predictions than the best program, T3SEpp. All our models performed well on the fungal data, with the two least-performing models predicting as well as WideEffHunter, the best-performing program. While we anticipated further improvements from model ensembles, our findings align with machine learning theory suggesting that ensemble benefits diminish with limited diversity among top-performing models and may inadvertently result in overfitting (Farhadi, Tatullo, and Ferrian 2025; Du et al. 2025).

The SHAP analysis showed that the contribution of features to the model performance varied: not all features contributed to the performance of XGBoost, whereas all features contributed to that of the Model_2. We also observed difference in ranking of features and in the magnitude of their contributions to the predictions. These differences could explain why Model_2 has a better performance over XGBoost. There is the limitation to consider that programs can incorrectly predict some proteins, which can affect the model’s final prediction. However, our models ultimately benefit from the improved performance that comes from integrating programs’ strengths.

We developed *fimep* (for integration of multiple effector prediction), a user-friendly Python package that incorporates the top-performing DNN Model_2. *fimep* offers prediction capabilities for individual kingdoms (bacteria, fungi, oomycetes) as well as cross-kingdom analysis. Our package allows researchers to preprocess outputs from all six evaluated programs, merge the processed data, encode the data, and generate reliable predictions through a single command-line interface or a step-by-step process. We have made the package, along with its usage and examples, publicly available on PyPI at https://pypi.org/project/fimep/.

## Conclusion

We combined the outputs of multiple effector prediction programs to develop models using both classical machine learning and deep learning approaches that accurately predict effectors from bacteria, fungi, and oomycetes than existing programs. By integrating predictions from several high-performing programs, we demonstrated that models built using this ensemble approach consistently outperform any single program. Unlike many existing tools that are limited to a specific taxon, our models can make accurate predictions across all three taxa: bacteria, fungi, and oomycetes. The tool we developed, called *fimep*, outperforms all the programs used in the study when evaluated on our test and validation data curated from all the programs.

## Acknowledgements

We wish to thank Josh Williams, Alison MacFadyen and George Deeks for help and discussion on machine learning, coding issues and HPC. We also wish to thank Norwich Bioscience Institutes Research Computing team for High Performance Computing support. The authors were supported by The Gatsby Charitable Foundation through a core grant to The Sainsbury Laboratory.

