## Supplemental Data for "Integrating and refining the predictions of ensembled plant effector detection programs using machine learning"

### Supplementary information

Table S1: Distribution of effectors and non-effectors data across training, test, and validation datasets

| Kingdom | Training data | | Test data | | Validation data | |
| --- | --- | --- | --- | --- | --- | --- |
|  | Effector | Non-effector | Effector | Non-effector | Effector | Non-effector |
| Bacteria | 149 | 150 | 260 | 260 | 259 | 259 |
| Fungi | 150 | 150 | 151 | 151 | 151 | 151 |
| Oomycete | 200 | 200 | 70 | 70 | 68 | 68 |

Table S2: Hyperparameter tuning ranges and values for each model type

| Models | Hyperparameter tuned | Values |
| --- | --- | --- |
| RF, XGBoost | n_estimators | 1 to 99 |
| RF, XGBoost | max_depth | None, 1 to 49 |
| RF | max_features | Sqrt, log2, None |
| RF | max_leaf_nodes | None, 1 to 49 |
| XGBoost | learning_rate | 0.1, 0.2, 0.01, 0.02, 0.001, 0.002 |
| XGBoost | subsample | 0.5, 0.7, 1 |
| SVM | C_range | 100 logarithmically spaced values between 0.01 and 100 |
| SVM | gamma_range | 100 logarithmically spaced values between 0.01 and 100 |
| SVM | kernel | rbf, linear |
| DNN | Number of layers | 1, 2, 3 |
| DNN | Number of neurons in the 1st layer | 16, 24, 32 |
| DNN | Number of neurons in the 2nd layer | 16, 32, 64 |
| DNN | Number of neurons in the 3rd layer | 16, 32, 64 |
| DNN | Regularizers | None, L1, L2 |
| DNN | Optimizers | Adam, SGD, RMSProp, AdamW |
| DNN | Learning rate for optimizers | 0.0001, 0.001, 0.01, 0.03162278, 0.1 |
| DNN | Learning rate for regularizers | 0.0001, 0.001, 0.01, 0.03162278, 0.1 |
| DNN | Dropout | 0.1, 0.2, 0.3, 0.4, 0.5 |

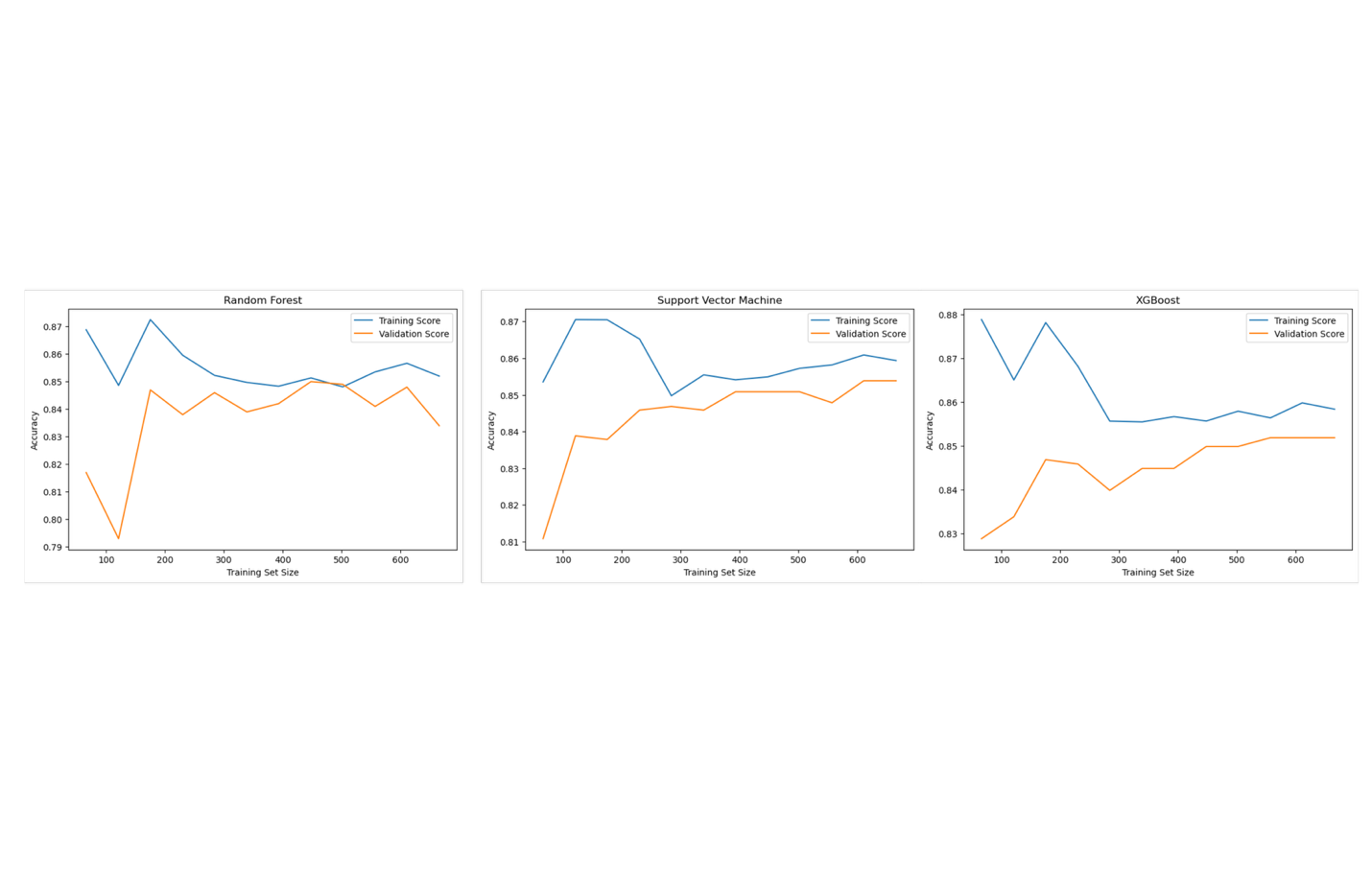

(a) Machine Learning Models

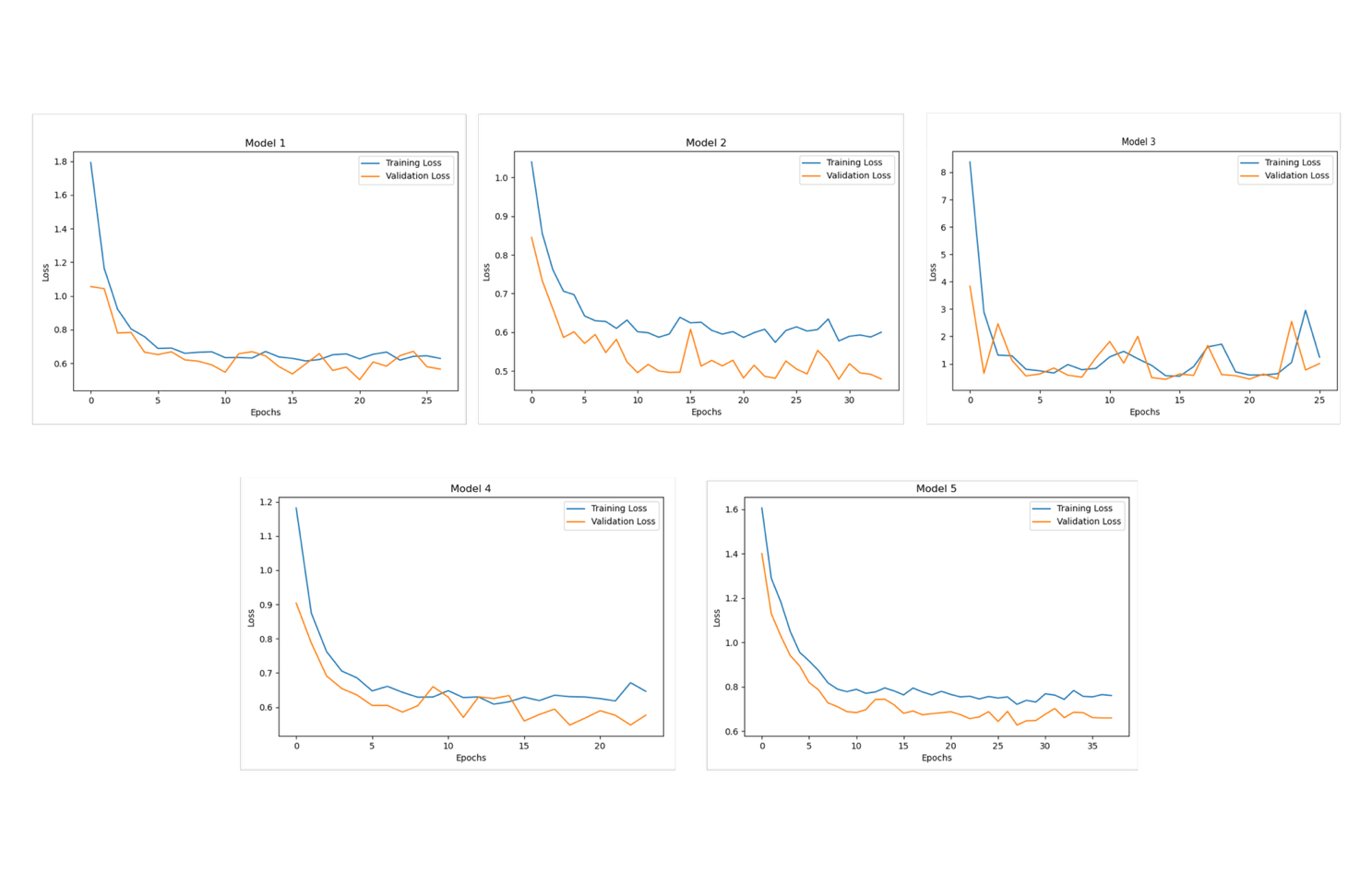

(b) Deep Neural Networks

*Figure S1: Learning curve for classical and deep learning models.* ***(A)*** *Learning curves of the training and validation accuracy plotted against increasing training set size for classical machine learning models (Random Forest, Support Vector Machine, and XGBoost).* ***(B)*** *Training and validation curve for the deep neural networks. The curve shows the reduction and convergence of the loss as the training progresses. The deep neural network models (Model_1 to Model_5)*

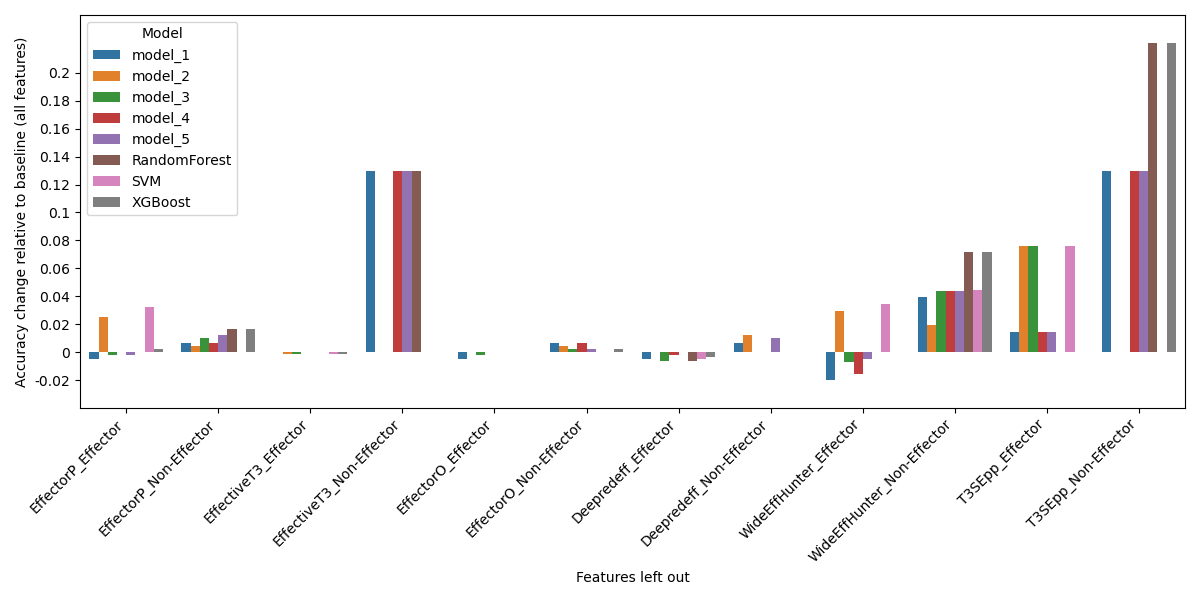

*Figure S2: Performance of the models when a feature is left out. The plot shows the features left out on the x-axis and the variation in accuracy from the baseline (when all models are present). XGBoost, extreme gradient boosting; RF, Random Forest; SVM, Support Vector Machine; the deep neural network models (Model_1 to Model_5)*
